# A guide for Single Particle Tracking: from sample preparation and image acquisition to the analysis of individual trajectories

**DOI:** 10.1101/2025.04.03.647105

**Authors:** Asaki Kobayashi, Junwoo Park, Judith Miné-Hattab, Fabiola García Fernández

## Abstract

Live-cell Single-Particle Tracking (SPT) is a powerful super resolution microscopy approach. It allows us to observe live recordings of individual molecules in living cells at high temporal and spatial resolution (100 Hz, 20 nm). Here, we discuss the implementation and application of super-resolution microscopy to quantify the mobility of single molecules in living human cells. We first describe the human cell sample preparation for SPT with an experimental time line overview, then super-resolution microscope setting and procedure to acquire SPT films for further SPT video analyses. We provide a practical guide to determine the quality of acquired SPT films in particular, number of photons and fluorescent background. Furthermore, we introduce several SPT video analysis software comparing their advantages and limitations. Expectantly, this SPT protocol will serve as a foundation for a better SPT performance.

## 1. Introduction

The way proteins diffuse and collectively interact plays an essential role for the good functioning of the cell. The mode of diffusion of a molecule inside the cell dramatically influences its interactions with the surrounding molecules [1]. Thus, investigating how molecules explore their environment and how their diffusion is affected in different biological contexts is crucial to understand biological processes. In optical microscopy, the spatial resolution is limited to 0.6 λ/NA ∼250 nm (where λ is the wavelength used to observe the sample and NA the numerical aperture of the objective). As a consequence, each fluorescent molecule appears as a spot of ∼250 nm in radius, and spots closer than this distance cannot be resolved. During the last 20 years, several techniques, referred to as super resolution microscopy, have emerged to break the diffraction limit allowing the imaging of structures smaller than 250 nm [2]. Among the different super resolution approaches, Single Molecule Localization Microscopy (SMLM) allows the visualization of single molecules. The key is to light only a small fraction of the fluorophores or dyes in the sample, so that their relative distance is greater than 250 nm. In this case, each spot in the image corresponds to a single molecule that can be localized at high spatial resolution by determining the centre of its point spread function (PSF), a common tool to describe the response of an imaging system to a point source. To obtain a full image of the sample, a few molecules are lighted on, localized and photobleached: this process is repeated for each frame of a movie.

SMLM approaches regroup three techniques: Photo Activation Localization Microscopy (PALM, [3], STochastic Optical Reconstruction Microscopy (STORM, [4]) and Single Particle Tracking (SPT, [5]). PALM and STORM are performed on fixed sample, to access the structure of macro-molecular complexes while SPT is performed in living cells to access the dynamics of individual molecules. Several other microscopy techniques can measure molecular dynamics in living cells (*e*.*g*., FRAP, FILM [6] or Hi-D [7]), however they give access to ensemble measurements (*i*.*e*., averaged diffusion coefficient, time and speed of fluorescence decay, residual fraction corresponding to the remaining fluorescence stably present after photobleaching). Unlike these approaches, SPT emerges as the best choice to measure the dynamics of individual molecules at high resolution. Today, the combination of new microscopy techniques, high-performance biological probes and image processing make SMLM a very accessible and standardized techniques in laboratories. Here, we present a workflow of the SPT approach, from preparation of cell samples, SPT acquisition to the analysis of molecular diffusion.

## 2. Sample preparation

In this method, human cell lines expressing a Halo-Tagged [8] were used. Cellular samples should be made using freshly prepared medium with appropriate antibiotics in an L2 laboratory. For analysis, cells in early passages (passages 3 to 8) are preferred and should be stored at 37°C in the dark.

### 2.1. Materials

1. Fresh culture cell media.
  - Growth media, DMEM media containing 10% FBS serum (Fetal Bovine Serum), glutamine (1%) and 1% antibiotics (Penicillin/Streptomycin).
  - Imaging media, L-15 (Leibovitz) media containing serum (20%), glutamine (1%) and antibiotics (1%).
2. Phosphate-buffered saline (PBS) solution (1x, pH 7.4).
3. Round cover glass #1.5, 25 mm (WWR 631-0172)
4. Cell chamber 35mm Attofluor (ThermoFischer)
5. Small forceps or needle
6. Sterile 6-well plates
7. Janellia Fluor dyes previously diluted in DMSO (stocked at 100 µM and several tubes of 1µM). Alternatively, other dyes can be used or cell lines expressing photo-activable fluorophores can be used [9]. Stable and endogenous tagged are better than transient transfection for the reproducibility of the results.

### 2.2. Methods

1. Day -1: Prepare a sterile 6-well plate. Place one coverslip in each well. Prepare at least two samples per condition. The number of coverslips depends on the number of samples. Harvest confluent cells (approx. 0.5 x 10^6^) to inoculate each well for 24h (**Figure 1**, step1).
2. Day 0: prepare a diluted solution of JF dye at 10 nM in PBS from the 1µM-DMSO stock.
3. Put 2 ml of diluted JF in PBS per sample. Shake carefully and leave for 30 minutes in a 37°C-incubator (**Figure 1**, step2).
4. During the resting time, prepare fresh imaging medium and maintain at 37°C.
5. Perform 3 washes with 2 ml of PBS for 10 minutes, removing the previous medium or PBS.
6. After the third wash, replace the PBS with 2 ml of imaging medium and provide an aliquot of few milliliters of imaging medium (1 ml per sample) into the microscope room.
7. Use a needle and forceps to carefully remove the cover glass from the 6 well plate to the cell chamber.
8. Proper assembly of the cell chamber (**Figure 1**, steps 3 and 4):
  - Place the coverslip on the round part of the cell chamber base with the help of the forceps.
  - Place the top of the chamber and close it carefully, without squeezing.
  - Add 1 ml of imaging medium and check that the medium does not leak out.
  - Place the assembly into a 37°C chamber. TOKAI system for example is compatible with all SPT microscopes, but an incubation chamber can also be used.

**Figure 1.**
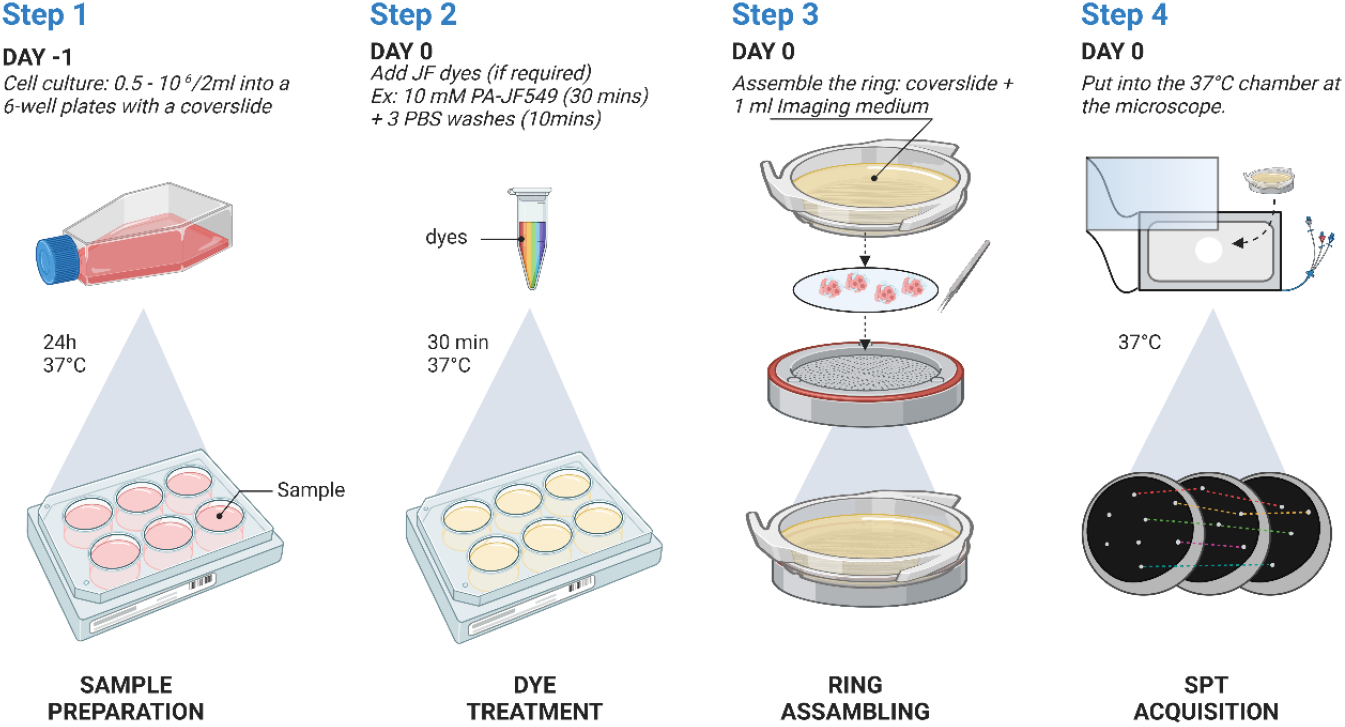
Sequential steps of sample preparation for SPT acquisition. One day prior to acquisition, cells should be harvested to be sufficiently confluent in sterile 6-well plates containing one coverslip per well. On the day of acquisition, cells are treated with the selected dye. In this example, cells are treated with PA-JF 549 for 30 min after 3 PBS-washes and final addition of imaging medium. Once the samples are prepared, coverslips are carefully placed into the cell chamber, imaging medium is added and samples are placed under the microscope, previously turned on and set up.

#### Note 2-1

If any drug treatment is planned during the experiment, it is recommended that it is added prior to the dye addition (Step 2).

## 3. Single Particle Tracking

Individual dynamic properties of single molecules and molecular interactions can be characterized using SPT. Analyzing their dynamics provides a better understanding of the molecular mechanisms in living biological systems. These approaches enable the acquisition of time-resolved trajectories from individual molecules which can be further analyzed using image processing programs.

In this section, a standard fluorescence-based SPT procedure to measure molecular motion of a single molecule in living human cells is described (see **Note 3-1**).

### 3.1. Materials

Materials should be equilibrated at room temperature, preferentially in a dark room or in a space partitioned with a blackout curtain to avoid light leakage, thus preventing fluorophore photobleaching.

#### 3.1.1. Super-resolution microscope

Presentation of the super-resolution microscope instrument as pictured in **Figure 2**:

**Figure 2.**
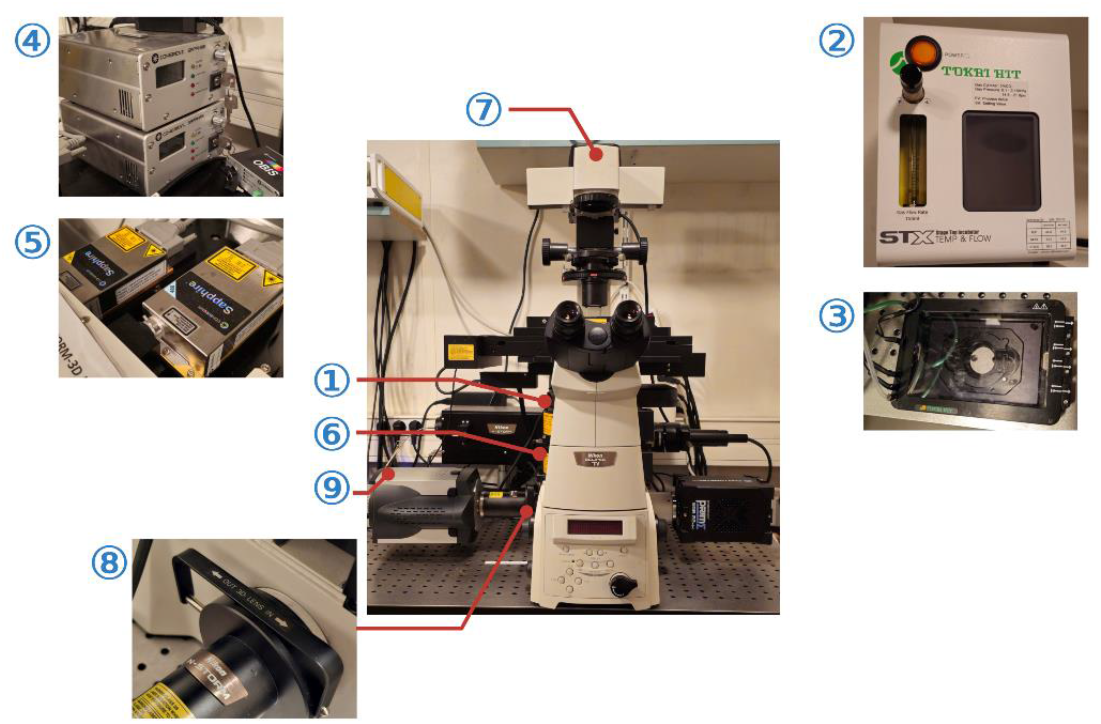
Pictures of the super-resolution microscope instrument used in this method. Numbers corresponds to the **section 3.1.1**.

1. 100 x oil immersion objective Nikon, numerical aperture (NA) 1.4 or more (Nikon)
2. Temperature controlling system (Stage Top Incubator, Tokai Hit for example)
3. Temperature controlling chamber (Tokai Hit for example)
4. Lasers controller
5. Lasers. Common lasers for SPT are 300 mW (or more) 561 nm and 640 nm, and 405 nm laser for the photo-activation. For single molecule imaging, the laser power at the objective is usually 1kW/cm^2^ or more.
6. Filter sets (Single band / Quad band) (**Note 3-2**). For a better signal to noise ratio, use single band emission filters.
7. Microscope main body with a perfect focus system (PFS) with a Highly Inclined and Laminated Optical sheet (HILO) excitation arm [10]
8. Astigmatism selection bar for 3 dimensional acquisitions
9. Camera EMCCD camera (iXon Ultra, ANDOR, or Hamamatsu Orca Fusion, or other models with similar quantum efficiency)
10. Computer with enough storage for super-resolution microscopic data (*e*.*g*., One SPT film of 500 frames with 10 illumination sequences corresponding to 5,000 frames approximately corresponds to 200 MB computer storage)
11. Acquisition software for SPT (from a company or home-made)
12. Immersion oil type F for 30°C (Zeiss)

### 3.2. Methods

The SPT data acquisition method described in this protocol relies on the imaging of human cell lines expressing a protein of interest fused to a HaloTag® [8], in conjunction with cell permeable dyes (Fluorescent dye *e*.*g*., Janelia Fluor 549 [8]). Human cell samples are kept within an incubation chamber at 37°C (with, if required, 5% CO^2^) in the dark during data acquisition.

1. Turn on the microscope and lasers one hour before the imaging to stabilize the laser (**Figure 2**).
2. Fill with high purity water the humidification canal in the temperature controlling system (Tokai Hit) and turn on the temperature controlling system (**Figure 2-2** and **2-3**) to stabilize the temperature in the chamber at 37°C (see **Note 3-3**).
3. Install the temperature controlling chamber on the microscope.
4. Prepared samples (*e*.*g*., human cells) in the metal cell chamber (see **Section 1**) can be loaded onto the microscope within the temperature controlling chamber.
5. From the centered laser position (calibrated laser position), adjust the laser angle for the Highly Inclined and Laminated Optical sheet (HILO) setting [10] (see **Note 3-4**). Activation of Janelia Fluor dyes (*e*.*g*., Photo-Activable Janelia Fluor 549® (HaloTag PA-JF549)) by the laser with 405 nm can help to figure out the right background/sample contrast. Once the laser angle is set, keep the same angle throughout the experiment during the day. Recalibration of the laser is required when turning on the microscope.
6. To keep the right focus on the sample (*e*.*g*., human nuclei), turn on the z-stack stabilizer function (Perfect Focus Systeme) (see **Note 3-5**). The switch is located on the microscope body (**Figure 2-7**).
7. Before starting the imaging, initial laser power needs to be set (see **Note 3-6**). Keep the total laser power as 100% in the laser power controller (**Figure 2-4**). Laser power which will be applied to the sample can be adjusted from the laser plane during the experiments.
8. Prior to SPT experiments, background fluorescence validation (see **Note 3-7**) is crucial. Apply higher laser power to single particle activatable level, then verify that the sample is not highly auto-fluorescent.
9. Laser 405 nm is used to activate the PA-JF549 fluorescent dye attached to HaloTag. The laser strength should be adjusted based on your samples. To track the “single’’ particles, adequate laser power, which will activate only a few molecules (not all molecules) should be determined (**Figure 3**). A continuous and very low 405 photo-activation should be used to start, *e*.*g*., 0.1% up to 1% depending on your sample. During the acquisition, the laser power can be increased upon signal breaching (see **Note 3-8**). For very abundant proteins, if the density of spots is too large, it is possible to use a pulsed 405 photo-activation, with for example 1 photo-activation every 10 images.
10. A maximal 1024 × 1024 pixels image (the final image size) can be captured, however focusing on a smaller area (*e*.*g*., a single nuclei) reduces image acquisition time leading to the protection of the sample from photo-toxicity. Use of the Region Of Interest (ROI) function to reduce the imaging area is highly recommended.

**Figure 3.**
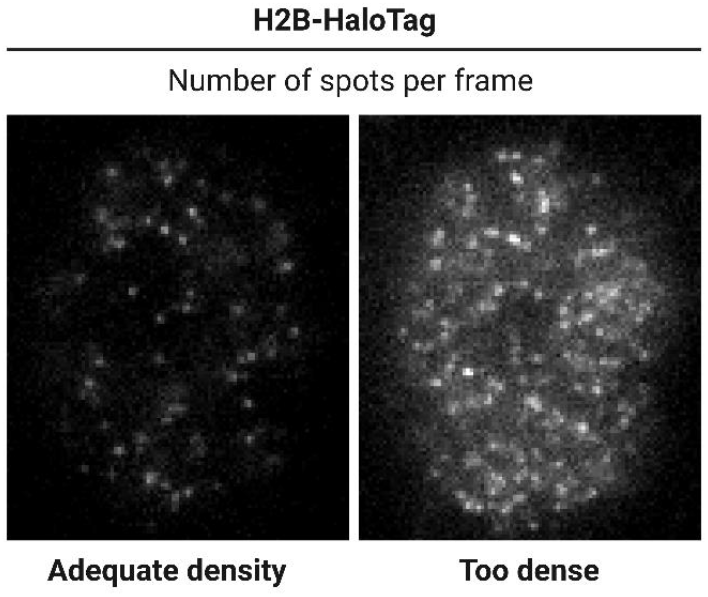
Examples of SPT films with an adequate number of spots per frames. On the right, U2OS cells expressing Halo-Tagged H2B bound to PT-JF549 are activated using a 405 nm laser. Weak JF activation (left) enables the acquisition of an adequate number of spots per frame to track single particle, while strong JF activation (right) shows too many spots per frame to track an individual particle. Too much JF activation per frame leads not only to photo-bleaching but also to spot miss-linking for further SPT analyses.

### 3.3. Notes

#### Note 3-1

For a detailed review about alternative scattering-based SPT method relying on non-fluorescent probes (*e*.*g*., plasmonic nanostructures) see [11].

#### Note 3-2

A single-band fluorescence filter set is installed in the microscope used in this method because it demonstrates higher performance by achieving high contrast and high signal-to-noise ratio images. Alternatively, quad-bands filter sets can also be used. Importantly, the choice of a filter set should be oriented toward minimization of the transmission gap to the sample in order to increase (and maximize) the sample/background contrast (see also **Note 3-3**).

#### Note 3-3

It is recommended to work with the same sample for a maximum of one hour. If data acquisition takes more than a few hours, supplementation of 5% CO^2^ is recommended for human cell samples.

#### Note 3-4

Among the super resolution microscopy techniques [12], conventional wide-field microscopy at high-resolution is best suited for thin samples. In contrast, thick samples, such as human cells, require more technical calibration because of the poor contrast due to the background fluorescence in a standard Köhler illumination setup. An adequate signal/background ratio is important for a higher resolution image which can be improved by using highly inclined and laminated optical sheet (HILO) microscopy [10]. In this case, the microscope setup is the same as a standard wide-field epi-fluorescence microscope as indicated in **section 3.1.1**. A key difference between the conventional microscopy and the HILO microscopy is the laser path setup. HILO microscopy employs an incident laser beam which is highly inclined by a large reflection and is laminated as a thin optical sheet at the specimen side resulting in a minimal illumination area, which is optimal for the speckle illumination of the sample with a higher resolution [13]. Indeed, HILO microscopy relies on the statistics of the speckle illumination pattern, thus it is noteworthy that protein diffusion inside the speckle plays a critical role in illumination contrast. Static diffusion exhibits high contrast and highly dynamic diffusion displays low contrast resulting in blurred image [14]. Characteristic illumination beam passage through the center of the specimen plane, always in a *z*-directional shift, makes the HILO microscopy successful at three-dimensional (3D) image acquisition. A *z*-dependent astigmatism can be introduced into the 3D images through the use of a weak cylindrical lens (see **section 3.1.1, Figure 2-8**) in the imaging path [15]. Point spread function (PSF) in 2D image presents a round shape where the PSF has equal widths in the *x* and *y* direction. Introduction of the cylindrical lens distorts the PSF into an elliptical shape where the fluorophore can be above or below the average focal plane. Alternatively, the image can appear as an ellipsoidal shape with its long axis along the *x* or *y*-axis.

#### Note 3-5

The Nikon Ti0-E inverted microscope series (see **section 3.1.1**) includes the Nikon Perfect Focus System (PFS) hardware component. The PFS controls axial movement of the microscope with a 5 millisecond (200 Hz) sampling rate which is independent from the microscope and camera controlling software. During the imaging, it is important to be aware of the possible artefacts resulting from drifts such as focus, mechanical or thermal drifts (*e*.*g*., due to physical, *z-axis* or *x-*axis and *y*-axis or laser heat, respectively). The simple focus/mechanical drift due to the axial movement can be corrected manually by subtracting the *x*/*y* and the *z* position of one of the reference beads (*e*.*g*., TetraSpeck Fluorescent Microspheres Sample Kit (Thermo Fisher Scientific)) from the selected traces. The thermal drift can be avoided by ensuring that temperature of the mechanical components (*e*.*g*., lasers) and ambient temperature are stable. Capturing the dynamic motion of single molecules in a single cell/nucleus requires a stable outline without these drifts. It is noteworthy that the drift can be verified by replaying the film backwards at the end of SPT film acquisition. In addition, it is critical to consider molecular blur motion, to adjust acquisition speed and to apply proper intervals in order to optimize and refine the imaging of live cells.

#### Note 3-6

The optical resolution increases at the expense of more exposure, longer acquisition times and or higher energy loads. Hence, while the optical resolution increases, photobleaching and phototoxicity also increase as a result [16]. A good balance (trade-offs) between expense of exposure time/energy load and photobleaching/phototoxicity need to be experimentally explored for an adequate higher resolution imaging. Illumination contrast plays a crucial role in higher resolution imaging (see also **Note 3-4**). Adjust laser input to generate sufficient illumination contrast between specific and unspecific photons in the sample.

#### Note 3-7

Cells and tissues are often auto-fluorescent, which is caused by natural light emission from its molecular components. Fluorescence background validation is important for an efficient capture of your fluorescence-emitting objects of interest. In this method we employ the HaloTag® system which offers a high performance in specific labelling. Resulting in an increase in background/sample contrast leading to higher resolution and decrease of false positive generation. Other tagging systems such as antibody labelling, endogenously tagged protein fusion *etc*., could also be considered. In any tagging system, it is crucial to validate auto-fluorescence prior to your experiences. Conventional microscopy can be used for the first sample quality test to eliminate possibilities of contrast issues from the higher resolution microscope. If background/sample contrast is still low in your system, reconsidering the imaging medium might be an option. In this protocol imaging medium without phenol red is used to avoid potential auto-fluorescence (see **Section 2**). In addition, intense washing steps of the imaging sample might also help in improving contrast “good enough” for photon detection for single molecule tracking.

#### Note 3-8

Super resolution microscopy is an optical microscopy technique that is advantageous for sample preservation. However, there are still some limitations regarding low photostability for long scales of time single molecule tracking (see also **Note 3-6**). Balancing the laser power to obtain good contrast and less phototoxicity/photobleaching during the single particle tracking is critical for a successful data acquisition. For that, minimising background auto-fluorescence to increase the sample-background contrast also helps. In this method, a fluorescent dye Janelia Fluor 549 (JF549) NHS ester (yellow fluorescent dye: excitation and emission maxima *λ*=549 nm and 571 nm) is used to attach to the HaloTag® system (see also **Note 3-7**). To eliminate false-positive fluorescent detection, a negative control (fluorescent dye in the non-tagged system) should be tested before starting the single particle tracking experiments. In addition, the selection of tags with good stability/biocompatibility and low cytotoxicity is important. Oversized tags with fluorescent dye may perturb the molecular dynamics in the living system. It is noteworthy that the plasmonic nanostructures are usually toxic to cells which further restrict their application in living cells (see also **Note 3-1**). Validation of protein function upon endogenous tagging should also be considered.

## 4. Analysis of SPT films

The microscopy video analysis of SPT experiments can be divided in two parts:

1. Inference of molecular trajectories from acquisitions.
2. Estimation of diffusive properties.

The trajectory inference consists of the localization of molecules from acquisitions and subsequent reconnections of detected molecules. Classical methods rely on the detection of high intensity peaks from the images and the search of the positions at pixel-level with local maxima. Recent methods based on machine learning use CNN (Convolutional neural network) [17, 18] were reported. In these approaches, the subsequent Gaussian fitting or centroid fitting approximates the positions of molecules at sub-pixel accuracy. Then, reconnection of localized molecules can be achieved using diverse methods such as nearest-neighbour or algorithms based on the combinatorial likelihoods [19, 20].

Reconstructed molecular trajectories can be thus used to estimate diffusive properties such as diffusion coefficients and diffusion exponent. The diffusion exponent [21] is a dependence where a molecular trajectory has an autocorrelation function for the consecutive movements over time. The MSD (Mean Squared Displacement, **equation 1**) is often used to estimate the diffusion coefficient when the autocorrelation is 0 (*i*.*e*., classical Brownian motion). However, estimation with MSD for anomalous diffusive molecules can lead to biased diffusion coefficient. Alternatively, the averaging of molecular positions with different time lags, TAMSD (Time Averaged Mean Squared Displacement, **equation 2**), and its averages on the ensemble (**equation 3**) provide statistical robustness for less-biased diffusion coefficient for sub-diffusive or super-diffusive molecules.

MSD at time *t* for a set of 1D trajectories can be calculated as follows:

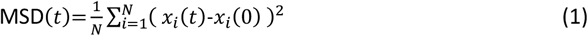

TAMSD for a trajectory of molecular *i* in 1D is defined as follows:

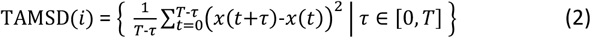

where *τ* is a time lag, and its ensemble-average for *N* molecular trajectories is as follows:

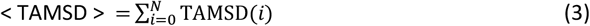

where < > denotes ensemble-average.

For accurate estimation, MSD based methods have been thoroughly scrutinized [22–24]. In the **Section 4-1**, some publicly available SPT software are introduced with a brief description. In the **Section 4-2**, we briefly describe the curves of ensemble-averaged TAMSD and squared displacement CDF (Cumulative Distribution Function) for 2 types of simulated trajectories as shown in the **Figure 4**. The experimental variety affecting the trajectory can be observed *via* displacements or mean jump-distances of trajectories. They are introduced in the subsequent section with simulated fBm (fractional Brownian motion) trajectories without considering experimental variety such as localization precision error, defocalization or blinking from microscopy.

**Figure 4.**
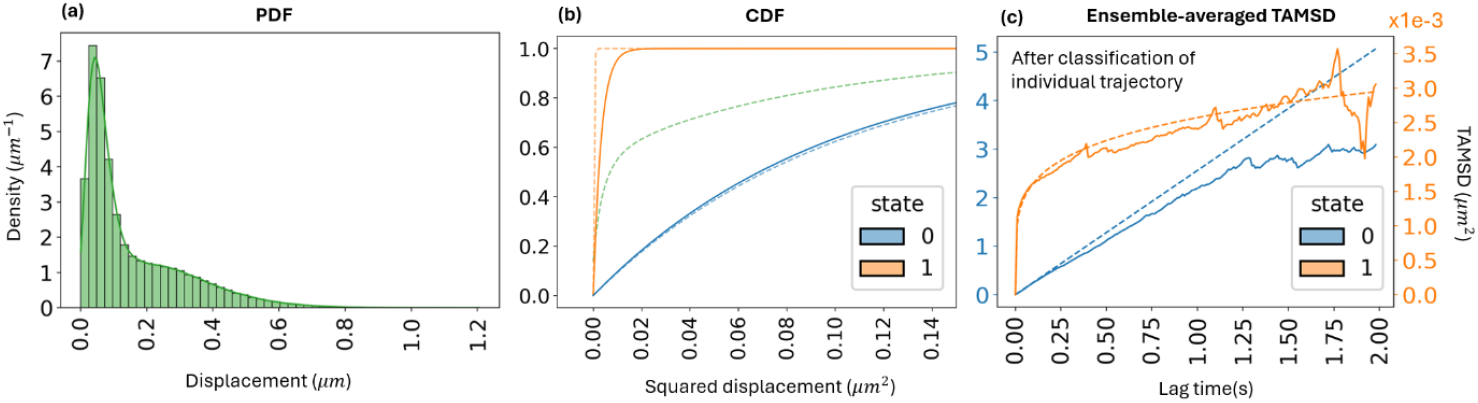
PDF of displacement with *τ*=10*ms*, CDF of squared displacement and ensemble-averaged TAMSD of two-state fBm trajectories. Trajectories are simulated with α=0.2, *K*=0.006 for a bounded (orange) state which is labelled as 1. And with α=1.0, *K*=2.56 for a free-diffusive (blue) state which is labelled as 0. (a) shows the displacements of all populations in green. Dashed lines are the ground-truths for each state. we can estimate the diffusion coefficient of two populations as in (b) without the categorical classification of trajectories at individual level. We can check the fitness of equation 2 in CDF. The fitting models show accurate estimation of diffusion coefficient for Brownian particles (blue curve, (b)), however it is less accurate for sub-diffusive molecules (orange curve, (b)). When a categorical classification of trajectories is performed for a given sample, we can also estimate the diffusive properties with ensemble-averaged TAMSD as shown in (c).

### 4.1. Inference of molecular trajectories from acquisitions

**Table 1** lists the most famous publicly available programs for SPT video prediction. For each program, we provide a brief description of the method and features of the software, including the availability of 2 or 3-dimensional tracking and the presence of an interactive GUI. We also highlight possible limitations in the method, such as the assumption of Brownian motion. Furthermore, some recently developed programs using neural network-based methods rely on a significant GPU processing capability for the full performance of the program in feasible computational time. Since there is no SPT program outperforming in overall aspects, the choice of program thus can be different depending on sample quality.

**Table 1.**
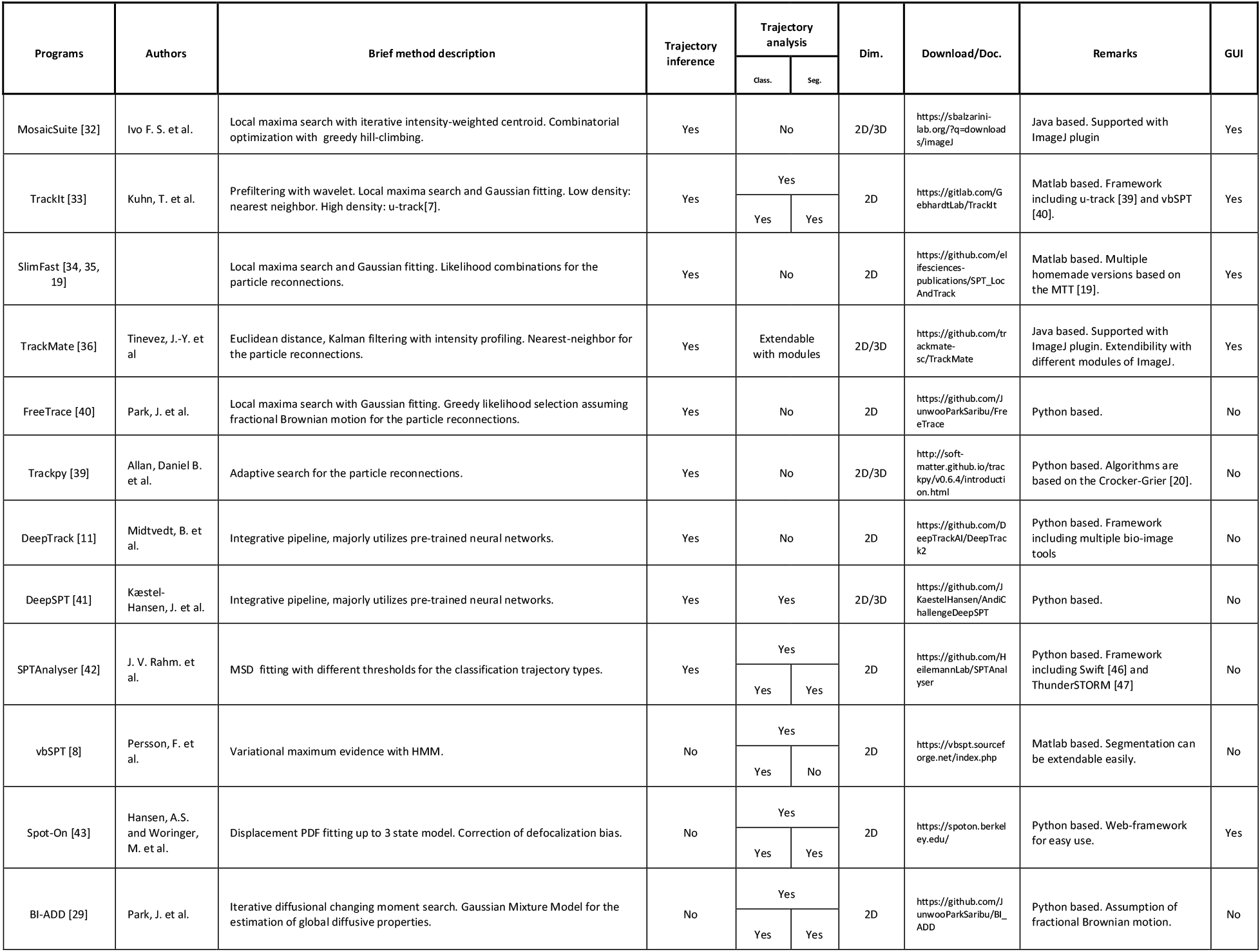
SPT software table. Only publicly available programs are listed. The following are the interpretations of column names: Trajectory inference: availability of the program to predict molecular trajectory from raw video. Trajectory analysis: availability of the program to estimate diffusive properties and classification and/or segmentation of heterogeneous trajectories. Class.: categorical classification of molecular trajectory. Seg.: segmentation of heterogeneous trajectory into multiple homogeneous. GUI: availability of graphical user interface for easy use.

The molecular density of video, signal-to-noise ratio, defocalization frequency followed by fast diffusion of molecules and blinking frequency of molecules are important factors for the choice of program. The number of input parameters are also an important feature for the minimal human-biased results. It is thus recommended to select the trajectory prediction program after inspecting the sample and the limitations associated with each program.

### 4.2. Estimation of diffusive properties

The measurement of diffusivity of molecules can divided in two categories of analysis:

1. Ensemble-level, where the quantification of diffusive properties is performed on a group of homogeneous trajectories. This approach is mainly used when experimental trajectories are too short to be analyzed independently.
2. Individual-level, where the quantification of diffusive properties is performed for each individual trajectory. This approach requires enough length of trajectory in general since low number of observation interrupt naturally the accurate estimate of diffusive properties.

Here, we will focus on the ensemble-level analysis, however some sophisticated or neural network based SPT software in **Table 1** also provide the diffusive property for each individual trajectory.

#### 4.2.1. Two methods to extract diffusion parameters at the ensemble-level

The first approach to quantify the mobility of molecules at ensemble-level is to build the CDF (Cumulative Distribution Function) which represents the cumulative of PDF (Probability Density Function). The CDF of displacements can be fit with the models described in **equation 4** or **equation 5** for different number of populations in a given sample. The example of PDF, CDF and ensemble averaged TAMSD is presented in the **Figure 4** for 2-population model [25–27]. Since the curve fitting can be solved using non-linear least square optimization for both PDF and CDF. However, the optimization with PDF is more complex than CDF due to numerical reason, it is recommended to fit the curve with CDF rather than PDF.

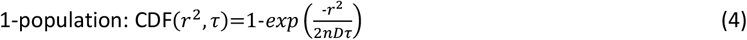

where *D* is the diffusion coefficient, n is the number of dimensions and *r*^2^ is the squared displacement for a time lag *τ*.

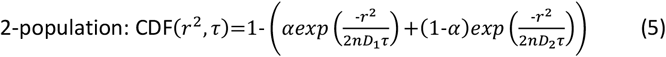

where *D*_*1*_ and *D*_*2*_ are the diffusion coefficients of sub-population 1 and 2 respectively, and α is the fraction of sub-population 1.

This method is adapted for data which cannot be analysed with MSD approaches due to the short length of trajectories. With the improvement of dyes and camera over the last 10 years, single molecules can now be visualized and tracked for longer times, improving considerably the quality of the data. In addition, the PDF/CDF method has several caveats. First, the approximation with CDF based on the Fick’s second law requires as a prior information the number of different populations in a given sample and sufficiently large sizes of sub-populations. Second, the model suits only classical Brownian particles and is not fit to anomalous diffusive particles. For anomalous motion, this approach gives an estimation of the diffusion coefficient but does not provide any information’s on the diffusive exponent, or the confinement radius for confined motion. Third, when the diffusion coefficient is slow (D < 0.1 μm^2^/s), the PDF/CDF approach is not precise enough and it is necessary to extract the diffusion coefficient from the experimental noise. In that case, it is recommended to use the MSD approach for the slow population (D < 0.1 μm^2^/s). For all these reasons, the PDF/CDF approach is less used.

When the trajectories are long enough, MSD approaches are better to extract diffusive parameters from SPT data. However, MSD based approach requires homogeneity of molecular trajectory to estimate the diffusive properties. In the large majority of the studies, the ensemble-averaged TAMSD is calculated (often referred as MSD) using **equation 3**. The curve can then be fitted to determine the best diffusion model: super-diffusive, Brownian or sub-diffusive (including anomalous and confined motion). A single trajectory can also exhibit several types of diffusion, with time, for example in the case of molecule transiently bound to a subtract, or exiting/entering a different compartment. Thus, in a heterogeneous medium, the trajectories need *a priori* segmentations for each diffusive type for accurate estimation since molecules change their dynamic over time. Some programs in the **Table 1** provide not only the categorical classification of trajectory types (classification of trajectory depending on the diffusive properties without segmentation in case of heterogeneous trajectory), but also the segmented trajectories (segmentation of heterogeneous trajectory into multiple homogeneous trajectories) and the estimated diffusive properties such as diffusion coefficient and diffusive exponent. After the classification of trajectories, if diffusive properties are not given by the programs, they can be measured by fitting the **equation 3** to the ensemble-averaged TAMSD of the molecular trajectories. The fitness of ensemble-averaged TAMSD is dependent on the number of long trajectory lengths. If the sample size is small with short trajectories, the fluctuation of ensemble-averaged TAMSD can lead to biased estimation results. It is thus recommended for large numbers of long trajectories.

The equation to estimate the diffusion properties of anomalous trajectories including classical Brownian motion is as follows:

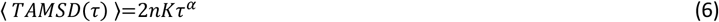

Where *n* is number of dimensions, *K* (*μm*^2^/sec^α^) is generalized diffusion coefficient, α is anomalous diffusion exponent, *τ* (*sec*) is lag time.

#### 4.2.2. Comparison on simulated trajectories

We simulated two types of 2,000 molecular trajectories [28] to compare briefly the different classical approaches to the ground-truth. One group follows sub-diffusive motion with α=0.2, *K*=0.006 in **equation 6**. Another group follows classical Brownian motion with α=1.0, *K*=2.56 which freely diffuses in 2D space. The simulated trajectories are classified in two classes: trajectories of slow molecules, likely corresponding to bounded molecules (orange) and trajectories of fast molecules, corresponding to free-diffusion (blue). All the trajectories before classification are also shown in green (**Figure 4** and **Figure 5)**. Figure 4 present the PDF, CDF and ensemble-average TAMSD for these simulations. If a molecule shows a classical Brownian motion, α is 1 and ensemble-averaged TAMSD is linear to elapsed time as shown as blue line in the **Figure 4C**. In longer time-scale, less trajectories can be observed in the data which fluctuates as shown in both bounded and free-diffusive motion. With CDF of squared displacement or ensemble-averaged TAMSD, we can estimate diffusive properties of molecules at ensemble-level for each population. The ensemble-averaged TAMSD shows its accurate estimation in many applications especially for the anomalous diffusive particles.

**Figure 5.**
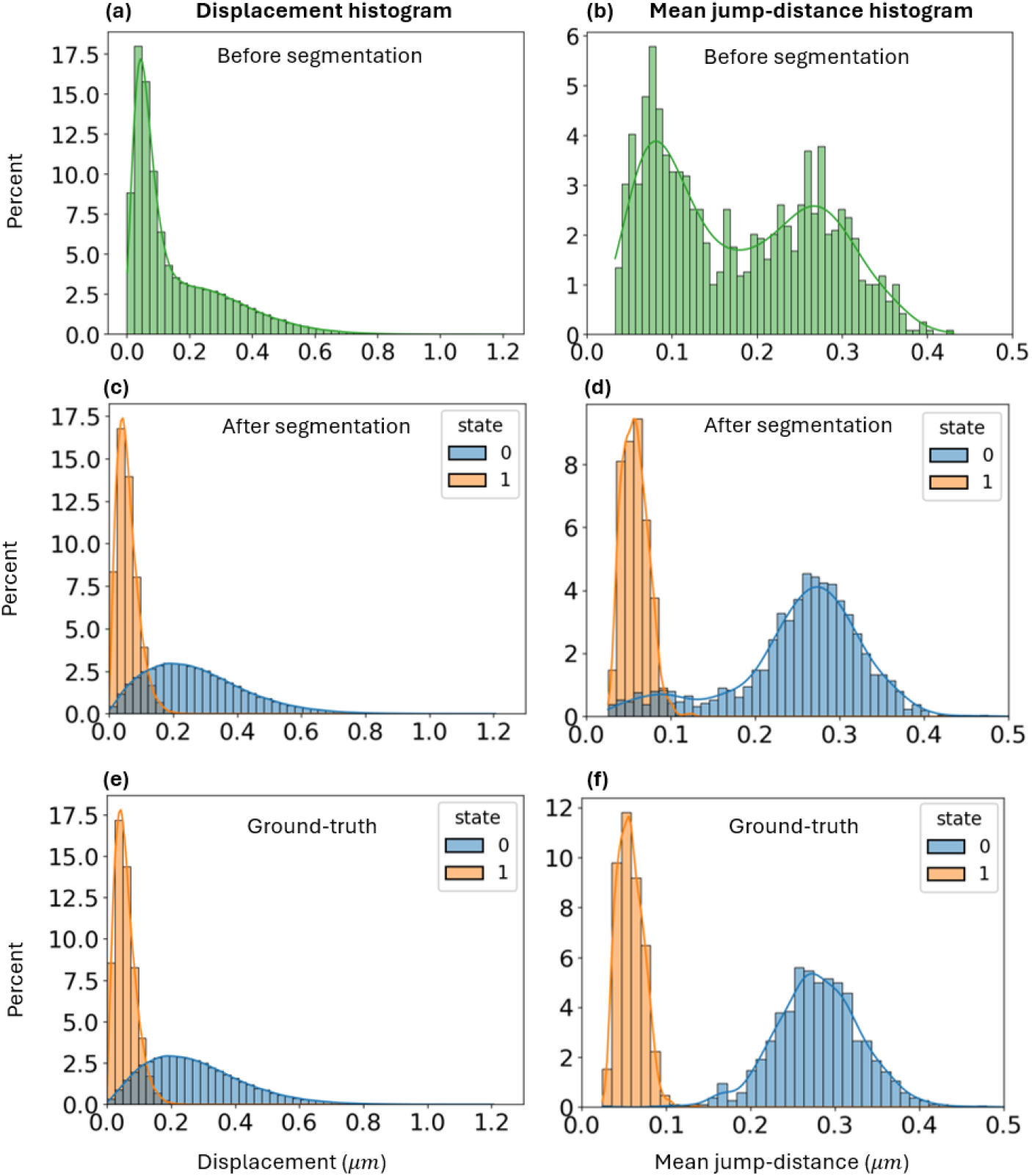
Displacement with *τ*=10*ms* and mean jump-distance histogram before and after segmentations of trajectories. The curves represent the continuity estimation of histogram for each state. The same simulated data and color codes are used as in Figure 4. The mean jump-distance histogram shows the importance of heterogeneous trajectory segmentation at individual level. (a) and (b) represent the data before segmentations of trajectories. (c) and (d) are segmented trajectories with BI-ADD [17]. (e) and (f) are ground-truth trajectories. The displacement histogram shows no influence induced by the heterogeneity of molecular dynamics. The effect such as irradiation on cells to quantify the difference of molecular dynamic [24], which induces a change of diffusive properties, can be quantified with Kolmogorov–Smirnov test by converting the displacement histogram to CDF. In the mean jump-distance, the effect is more directly observable, which can be seen through a shift in the distribution.

However, its use is limited for long and sufficiently large number of trajectories to ensure statistical robustness.

Another approach to quantify the molecular dynamic in scalar consists in observing the mean jump-distance (**equation 8**) of trajectories. The set of displacements for a given time interval *τ* and mean jump-distances for given trajectories in 1D are defined as follows:

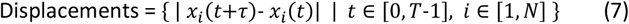

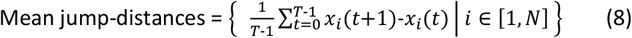

where *i* is an index of trajectory, *T* is the trajectory length.

This feature can provide additional information about the trajectory data. **Figure 5** shows the same data analysed with displacement and mean jump-distance histogram. We can see that the displacement histogram is not influenced by the heterogeneity of trajectories, and enables to estimate the diffusion coefficient of Brownian particles by converting it into CDF. In the mean jump-distance, the distinction between different populations can be clearer than in the displacement histogram. However, if the trajectories are not segmented for each type, this may lead to a biased result as we can see in the **Figure 4**. In the **Figure 5D** and **Figure 5F**, we can observe the effects of heterogeneity segmentation of trajectory in mean jump-distance. (c) and (d) are the result of a priori segmented trajectories with BI-ADD [29] and (e) and (f) are the ground-truths. The mean jump-distance shows Gaussian distribution according to CLT (Central Limit Theorem) assuming identical and independent distribution, which makes it easier to fit each population with Gaussian function. However, a small number of populations of molecule in a given system can be hidden underneath larger populations in both displacement and mean jump-distance histogram.

Finally, several studies use the distribution of angles for all trajectories to extract specific information from SPT [30]. If some molecules of a given sample show directed motion, the ensemble of relative angular differences of trajectory for a consecutive time interval can show the significance for both trajectory classification and subsequent analysis of the sample. However, this angular property of molecular trajectory can hardly have a significant meaning in many samples due to the weak dependence or independence of molecular diffusivity over time unless the molecule shows quasi-directed motion. In consequence, the statistical choice to analyse the trajectory data varies depending on the molecular diffusive properties of the sample.

Ensemble-level analysis can provide the diffusion coefficient for a group of trajectories. However, the analysis of trajectories at individual-level can be more informative with the identification of heterogeneity, but also more challenging. The visualization of identified molecular trajectories can also give the information such as local density of trajectory inside a cell or nucleus or difference of diffusivity of trajectory depending on a locus. The software InferenceMap [31] provides such visualizations for molecular trajectories and the diffusivity with GUI assuming the motion of molecules with an overdamped Langevin equation. In conclusion, the estimate of diffusion exponent at individual-level explaining the anomalous molecules is nearly intractable for short length of trajectories due to low number of observations. Recent neural network-based methods enlarge the possibility of this estimation at individual-level and can give us more information of molecular behaviour inside cell.

## 5. Concluding remarks

This guide provides instructions for both beginners and experienced users to perform SPT microscopy, from experiments to analysis. With the standardization of super resolution microscopes and the development of new AI-based methods to analyse the trajectories of single molecules, SPT is changing our view on cell biology. Today, the SPT approach plays a significant role in enhancing our understanding of fundamental biological processes. Importantly, by investigating the interactions of a proteins of interest *in vivo* with external reagents, it can also be applied on a large scale for drug discovery.

## 6. Acknowledgements

This work was supported by ANR-18-CE12-0015-03 (RepairChrom) and ANR-22-CE12-0039 (AROSE). The J.M-H team was financially supported by the i-Bio Initiative from the Idex Sorbonne University Alliance, the IBPS Incentive Action and the ATIP Avenir 2021. We are grateful to Adrien Bérard for his fruitful comments on the manuscript.

## Notes

### Competing Interest Statement

The authors have declared no competing interest.

